# Fast and accurate genome-wide predictions and structural modeling of protein-protein interactions using Galaxy

**DOI:** 10.1101/2021.03.17.435706

**Authors:** Aysam Guerler, Dannon Baker, Marius van den Beek, Bjoern Gruening, Dave Bouvier, Nate Coraor, Stephen D. Shank, Jordan D. Zehr, Michael C. Schatz, Anton Nekrutenko

**Affiliations:** Dept. of Computer Science, Johns Hopkins University, Baltimore, MD, USA; Dept. of Biochemistry and Molecular Biology, Penn State University, College Park, PA, USA; Dept. of Bioinformatics, Freiburg University, Freiburg, Germany; Institute for Genomics and Evolutionary Medicine, Temple University, Philadelphia, PA, USA

## Abstract

Protein-protein interactions play a crucial role in almost all cellular processes. Identifying interacting proteins reveals insight into living organisms and yields novel drug targets for disease treatment. Here, we present a publicly available, automated pipeline to predict genome-wide protein-protein interactions and produce high-quality multimeric structural models.

Application of our method to the Human and Yeast genomes yield protein-protein interaction networks similar in quality to common experimental methods. We identified and modeled Human proteins likely to interact with the papain-like protease of SARS-CoV2’s non-structural protein 3 (Nsp3). We also produced models of SARS-CoV2’s spike protein (S) interacting with myelin-oligodendrocyte glycoprotein receptor (MOG) and dipeptidyl peptidase-4 (DPP4). The presented method is capable of confidently identifying interactions while providing high-quality multimeric structural models for experimental validation.

The interactome modeling pipeline is available at usegalaxy.org and usegalaxy.eu.

## Introduction

Obtaining a complete map of interacting proteins is crucial to decipher the inner workings of living organisms. Among many other roles, proteins act in dynamic collaboration to fulfill biological functions by catalyzing chemical processes. Commonly, interactions are elucidated through a variety of experimental methods (Shoemaker, 2007, Zhou, 2016) which are capable of evaluating an ever-larger number of putative protein pairs. Unfortunately, the overlap between these methods is often limited which either indicates a high false positive rate or a low coverage. Often 40 to 90% of the detected interactions do not overlap between different methods (Mering, 2004, Rao, 2014). Also, high throughput methods do not provide structural insights into the formed protein-protein complex. More reliable methods such as crystallography and NMR spectroscopy do yield structural information but are labor intensive and as such only applicable to a limited number of proteins. In a recent study we demonstrated that the gap between low and high throughput methods can be bridged by identifying distantly related protein-protein homologues with similar protein-protein interfaces (Gong, 2021). Application of the SPRING method (Guerler, 2013) to *Escherichia coli* competitively identified protein-protein interactions while producing accurate multimeric protein structure models of which 39 by now have been confirmed in high-resolution experiments. Other studies applied our method to the minimal synthetic genome syn3.0 (Zhang, 2021) and the mouse genome (Li, 2016). In the present study we describe how we implemented our pipeline on Galaxy (Afgan, 2018), a web-based computational workbench used by many scientists across the world to analyze large data sets. This allows scientists to reproduce, share and embed the resulting interactome networks within their own analysis pipelines. Given a set of query sequences and a list of known protein structures, the pipeline employs SPRING with HHsearch (Steinegger, 2019), and TMalign (Zhang, 2005) to detect and structurally model protein-protein interactions. We validate the pipeline’s performance by comparing the resulting Human and Yeast protein networks with experimental findings. Similar to the results for *Escherichia coli*, the method competitively resolves Human and Yeast protein-protein interaction networks. As novel targets, we identified Human proteins likely to bind the papain-like protease of SARS-CoV2’s non-structural protein 3 (Nsp3). We also obtained models for SARS-CoV2’s spike protein (S) in complex formation with myelin-oligodendrocyte glycoprotein (MOG) and dipeptidyl peptidase-4 (DPP4). Some of the detected interactions have already been experimentally confirmed in recent literature, others provide novel insights into the pathology of SARS-CoV2. Notably, the interaction with DPP4 has been suggested to cause a higher mortality rate of diabetics contracting SARS-CoV2 (Valencia, 2020) while the MOG receptor is associated with the MOG antibody disease which relapses in SARS-CoV2 patients (Woodhall, 2020).

## Methods

### Protein-Protein Interaction Analysis Pipeline

We present a Galaxy pipeline to predict and structurally model protein-protein interactions on genomic scale. The pipeline takes the following inputs:

1. An individual file or a pair of files containing multiple FASTA entries of protein coding sequences. The pipeline will attempt to identify protein-protein interactions within the set of query sequences.
2. Text file containing the list of all Protein Data Bank (PDB) (Berman, 2000) entry identifiers to be employed as a multimeric template library. This step can be skipped if the library has already been constructed.
3. PDB70 threading library files as provided by the developers of HHsearch. These files are used to perform single-chain threading and can be obtained from http://www.user.gwdg.de/~compbiol/data/hhsuite/databases/hhsuite_dbs/.

The following outputs are generated:

1. Tabular file containing all identified interactions with their corresponding templates and Z_com_ scores (Wong, 2021).
2. Tabular file containing the details of the produced multimeric structural models and the corresponding model properties, i.e. SM-score, TM-score, a knowledge-based contact energy term E_contact_ and the fraction of inter chain clashes.
3. Collection containing dimeric structural models for each interaction, including the structures of the identified templates from the PDB.
4. Bar chart displaying the prediction accuracy in comparison to experimental results derived from the BioGRID database.

### Available Workflows

We build two analysis workflows, one for intra-genome and another for inter-genome protein-protein interaction prediction (see **Table 1**). The source code of the entire pipeline, including data preparation and interaction prediction logic is written in Python 3 and publicly available at https://github.com/guerler/springsuite under the GNU License. All methods can be executed locally or using the Galaxy web interface on usegalaxy.org and usegalaxy.eu.

**Table 1.**
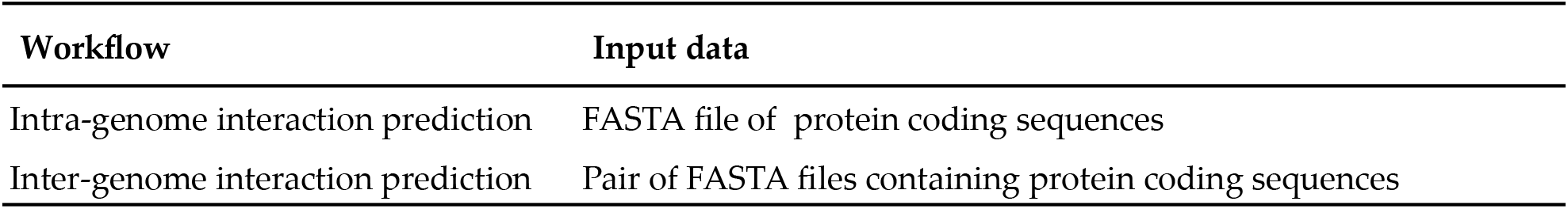
Galaxy workflows available at usegalaxy.org and usegalaxy.eu.

### Data Preparation

The presented pipeline utilizes all protein-protein interfaces available in the PDB as a template library for interface homology detection (**Figure 1**). The data preparation starts by using the DBkit tool in Galaxy to download all PDB entries and store them as a ffindex/ffdata database pair. As of November 29^th^, 2020 this amounted to 170,860 files. Then the SPRING Cross tool is applied which scans each PDB entry for protein-protein interfaces and stores the corresponding interacting PDB chain identifiers in a 2-column lookup table as a pairwise index of all interactions. In more detail, the SPRING Cross method proceeds by using the PDB REMARK 350 entries to build all bio units available in a given PDB entry. Then all C-alpha atom distances between two separate chains within the same bio unit are determined. If more than five distances below 10Å are detected for a pair of PDB chains, the corresponding PDB chain identifiers are deemed as interacting and added as a new row to the resulting 2-column lookup table. This yields a complete set of 988,784 interacting PDB chain identifier pairs contained in the PDB which we will use as a multimeric template library.

**Figure 1.**
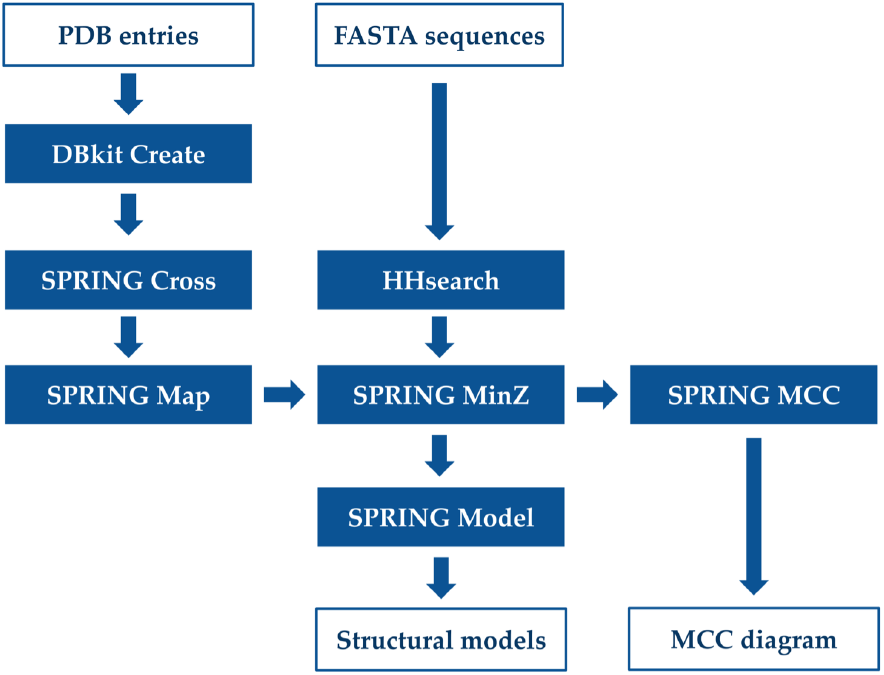
Schematic overview of the presented pipeline, illustrating the main input data sets, tools and outputs.

In a consecutive step the SPRING Map tool is applied which uses PSI-BLAST (Altschul, 1997) to detect close homologues of the PDB70 database for each PDB chain identifier listed in the columns of the lookup table. The identifiers of matching PDB70 entries are added in two additional columns to the lookup table. This allows us to apply HHsearch on a non-redundant subset of the PDB, containing entries with less than 70% sequence identity to each other. Although possible, expanding the monomeric threading database by including every PDB chain would significantly impact the database preparation time without improving the overall prediction performance. We used the PDB70 database issued on November 18, 2020 containing 58,900 entries. If a PSI-BLAST E-value equal to zero is used, 257,698 interaction frameworks which exactly match the sequences in the PDB70 are detected. With an E-value threshold of 0.001, the resulting 4-column lookup table contained 900,772 interaction frameworks suitable for the monomers available in the PDB70 database.

### Interaction Prediction

The pipeline’s interaction prediction logic uses SPRING with HHsearch and TMalign, and was designed to exploit the redundancy of available protein-protein interfaces in order to predict and model novel protein tertiary structures.

Initially each query sequence Q is threaded by HHsearch against the PDB70 monomeric template library to identify a set of putative templates (T_i_, i=1,2,…) each associated with a Z-score (Z_i_). The Z-score is defined as the number of standard deviations by which the raw alignment score differs from its mean. A higher Z-score indicates a higher significance and usually corresponds to a better alignment.

Considering all possible target sequence pairs, the SPRING Min-Z tool uses the previously described 4-column lookup table to select interaction frameworks which are shared by the monomeric templates of a query pair. The Z-score of the framework is defined as Z_com_ which is the smaller of the two monomeric Z-scores. A more detailed description of this algorithm is provided in Gong et al. 2021 and Guerler et al. 2013.

### Interaction Validation

The accuracy of predicted protein-protein interactions is evaluated using the SPRING MCC tool. This tool compares the set of interactions from SPRING with interactions obtained from experimental methods contained in the Biological General Repository for Interaction Data sets (BioGRID) (Oughtred, 2020). BioGRID is an open access database that contains protein interactions curated from primary biomedical literature for all major model organism species and Humans. The SPRING MCC tool accesses the BioGrid Tab 3.0 format columns 24 and 27, containing the UniProt (UniProt Consortium, 2021) accession identifiers of interacting protein pairs. The method only operates on interactions identified for sequences which are available in the UniProt database.

Initially, the SPRING MCC tool produces a ‘negative’ data set of non-interacting protein pairs by randomly sampling protein-protein interaction pairs from the set of query protein sequences. If a UniProt localization file is provided, the non-interacting pairs can be determined by sampling protein sequences from different subcellular regions. This approach can reduce the false-negative rate of the resulting negative data set.

Subsequently, the protein-protein interaction sets identified by each experimental method are considered to be truly interacting, constituting the ‘positive’ data sets for the cross-validation process. Each method is compared to all other methods using the positive data sets and a negative data set of equal size. The resulting Matthew’s correlation coefficients (MCC) are plotted using the Matplotlib library (Hunter, 2007). An example of such a plot is shown in **Figure 2**, displaying the results for the Human and Yeast interactome validation. The legend lists the experimental methods used to determine the corresponding positive data set.

**Figure 2.**
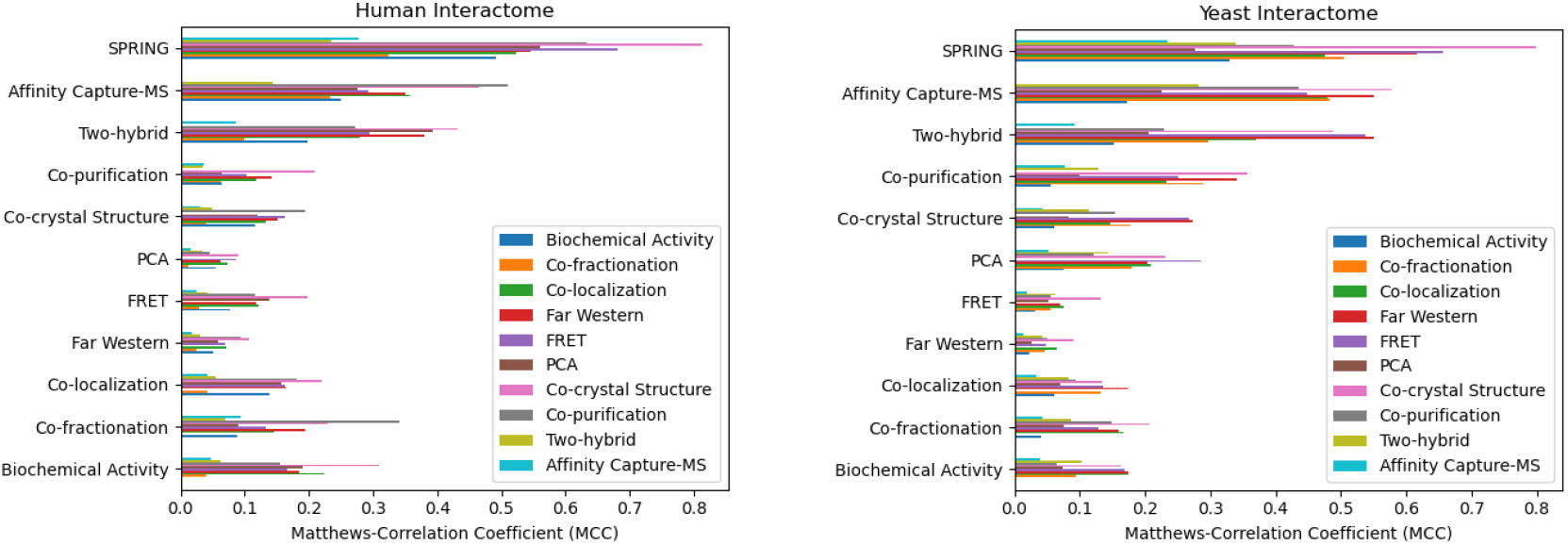
Human (left) and Yeast (right) protein-protein prediction results and comparison. Showing Matthew’s correlation coefficients (MCCs) as produced by comparing SPRING predictions with different protein-protein interaction experiments available in the BioGRID database. Each experimental method serves as a positive validation set for every other method.

### Structural Modeling

If a pair of proteins, Chain A and Chain B, is deemed to potentially interact e.g. Z_com_ > 25, the complex structure is constructed by structurally aligning the top-ranked monomer templates of Chain A and Chain B to all putative interacting frameworks using the SPRING Model tool which utilizes TM-align. The structural alignment is built on the subset of interface residues. The resulting models are evaluated by the recently established SPRING model score (Vangaveti, 2020):

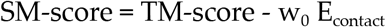

where TM-score is the smaller TM-score returned by TM-align when aligning the top-ranked monomer models of Chain A and Chain B to the interaction framework; E_contact_ is a residue-specific, atomic contact potential derived from 3,897 non-redundant structure interfaces from the PDB using the formula of RW (Zhang, 2010). The weight parameter w_0_ is set to 0.01 through a training set of protein complexes to maximize the modeling accuracy of the interface structures. The final model is evaluated for clashes and removed if more than 10% of the resulting C-alpha atom contacts share a distance of less than 5Å between the interacting pair of protein structures.

## Results

### Performance Validation with Human and Yeast Interactomes

The pipeline’s performance is validated on 20,610 raw protein coding gene sequences from the Human Reference Genome (UP000005640) of the UniProt database. This process evaluates ∼212 million possible pairs to identify the set of interacting protein-protein pairs. Each interaction is ranked by the Z_com_ score and Matthew’s correlation coefficient (MCC) is determined with regard to a negative data set of non-interacting protein pairs produced by the SPRING MCC tool and positive data sets derived from each experimental method. The negative data set has been sampled to contain proteins from different subcellular regions. **Figure 2** displays the cross-validation performance results in comparison to ten experimental methods available in the BioGRID database. Note that we applied SPRING on the raw protein coding sequences without separating the individual proteins using the CDS record provided by GenBank (Benson, 2014). In total the 20,610 protein coding genes encode for about 75,776 individual proteins.

We repeated the same experiment using the Yeast genome (UP000002311) to identify protein-protein interaction networks. In total 6,045 protein coding genes were parsed through the pipeline evaluating ∼18 million possible protein-protein interactions using the public Galaxy instance at https://usegalaxy.org. The results are shown in the right panel of **Figure 2**.

A more detailed analysis regarding the prediction performance of the presented pipeline versus experimental methods has been recently published for the *Escherichia coli* genome (Gong, 2021). The pipeline predicted several protein complex structures which were later experimentally verified by crystallography.

For all three genomes, our pipeline was able to implicitly identify individual protein sequences and achieve an overall performance which is comparable if not better than existing experimental methods.

### SARS-CoV2 protease (Nsp3) and Ubiquitin

We next applied the Galaxy pipeline to the genome of SARS-CoV2 which causes a novel severe acute respiratory syndrome and has been declared a pandemic (Naqvi, 2020). The SARS-CoV2 genome contains 13 to 15 open reading frames with ∼30 thousand nucleotides, including 11 protein-coding genes. Our pipeline identified Human substrates for the papain-like protease of SARS-CoV2 which is part of the non-structural protein 3 (Nsp3) (see **Fig. 3**).

**Figure 3.**
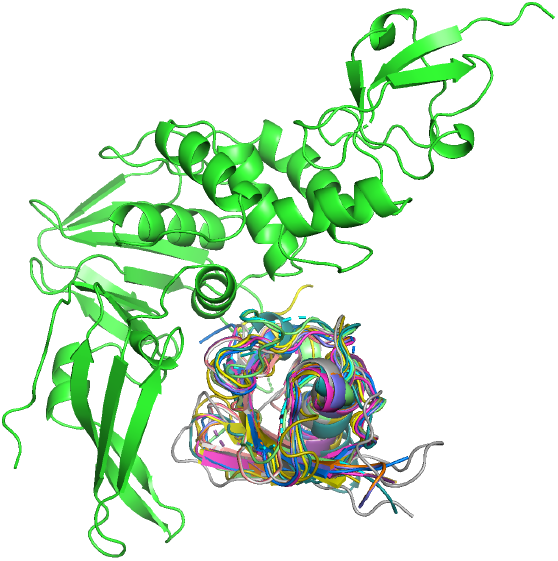
Putative ubiquitin-like substrates (colored) of SARS-CoV2 papain-like protease (green).

**Table 2** shows a list of the highest ranking fifteen substrates with matching multimeric templates and model quality attributes i.e. SM-score, TM-score, E_contact_ and Z_com_. The two highest ranking interactions were identified for ISG15 (SM-score=1.11) and ANKUB (SM-score=1.10). ISG15 has recently been experimentally confirmed as a substrate (Shin, 2020) and ANKUB was suggested in a computational cleavage enrichment study (Prescott, 2020).

**Table 2.**
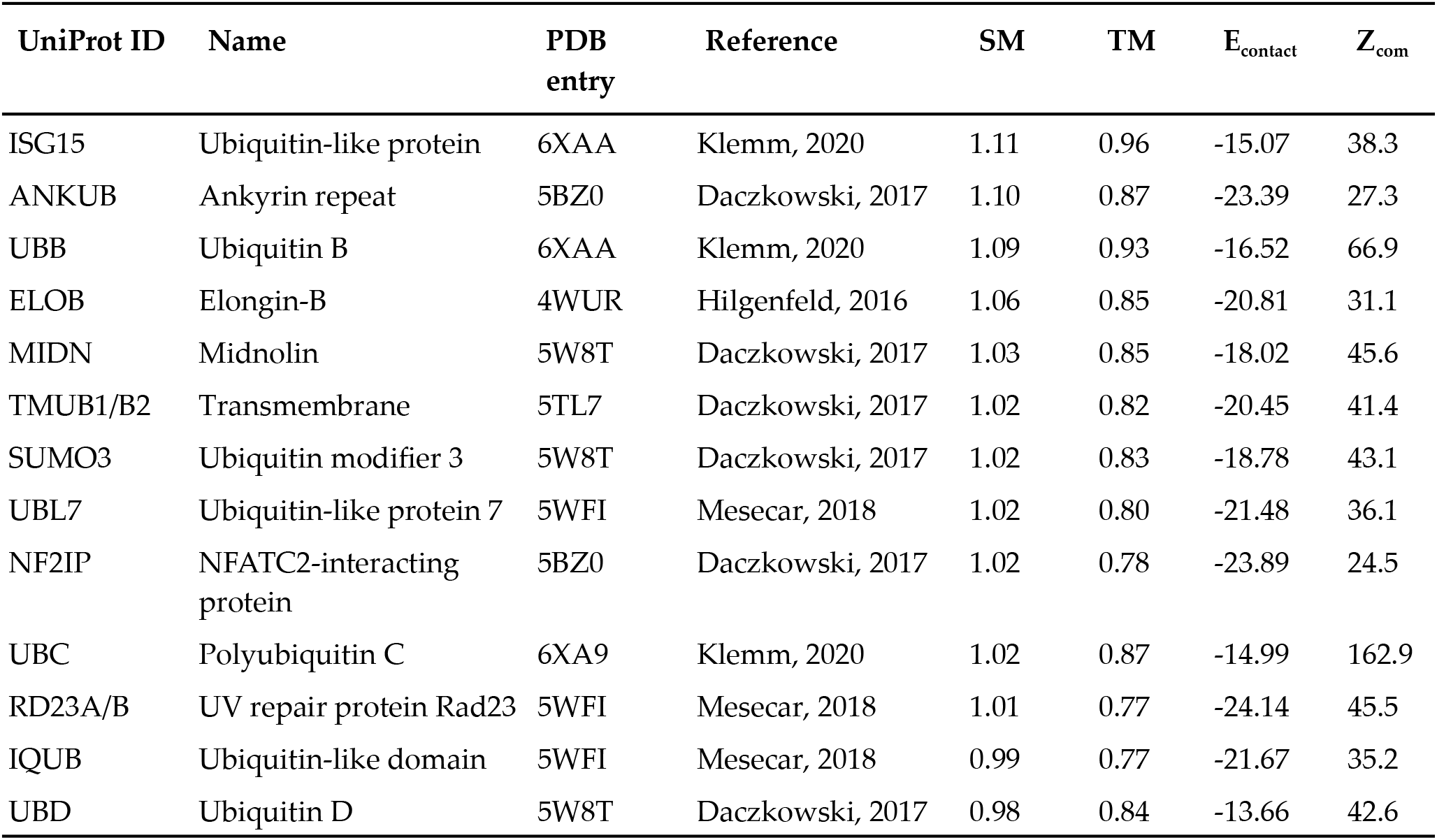
SARS-CoV2 papain-like protease substrates. Top scoring Human protein complex models for SARS-CoV2 papain-like protease of Nsp3.

### SARS-CoV2 Spike protein (S) and Myelin-oligodendrocyte Glycoprotein

We also identified Human proteins interacting with SARS-CoV2’s spike protein (S). The top-ranking interaction was found for angiotensin (ACE2/ACE), which is widely known to be the primary receptor for SARS-CoV2 (Peng, 2020, Zhou, 2020).

The second highest ranking model was detected for the interaction with the myelin-oligodendrocyte glycoprotein (MOG, see **Fig. 4**). MOG is a protein located on the surface of myelin sheaths in the central nervous system (Kezuka, 2018). Our pipeline modeled the monomeric structure of MOG with the highest ranking homologue in the PDB70 database which is PDB entry 4PFE (Eshaghi, 2015) at a Z-score of 102.2. We compared the resulting monomeric model with the model provided by Mesleh et al. in 2002. Both models resolve MOG as a beta-barrel and share significant similarity at a TM-score of 0.70. Additionally several suitable multimeric template frameworks were identified. The corresponding PDB entries are 7C8V (Li, 2020), 6XC2 (Yuan, 2020), 6XC4 (Yuan, 2020), 7BZ5 (Wu, 2020) and 7C01 (Shi, 2020). All of these structures, except 7C8V, were crystalized with a potent neutralizing antibody of SARS-CoV2. **Table 3** shows the identified template frameworks and the resulting model scores. The results indicate two distinct putative binding modes which may occur in tandem (see **Fig. 4C**).

**Figure 4.**
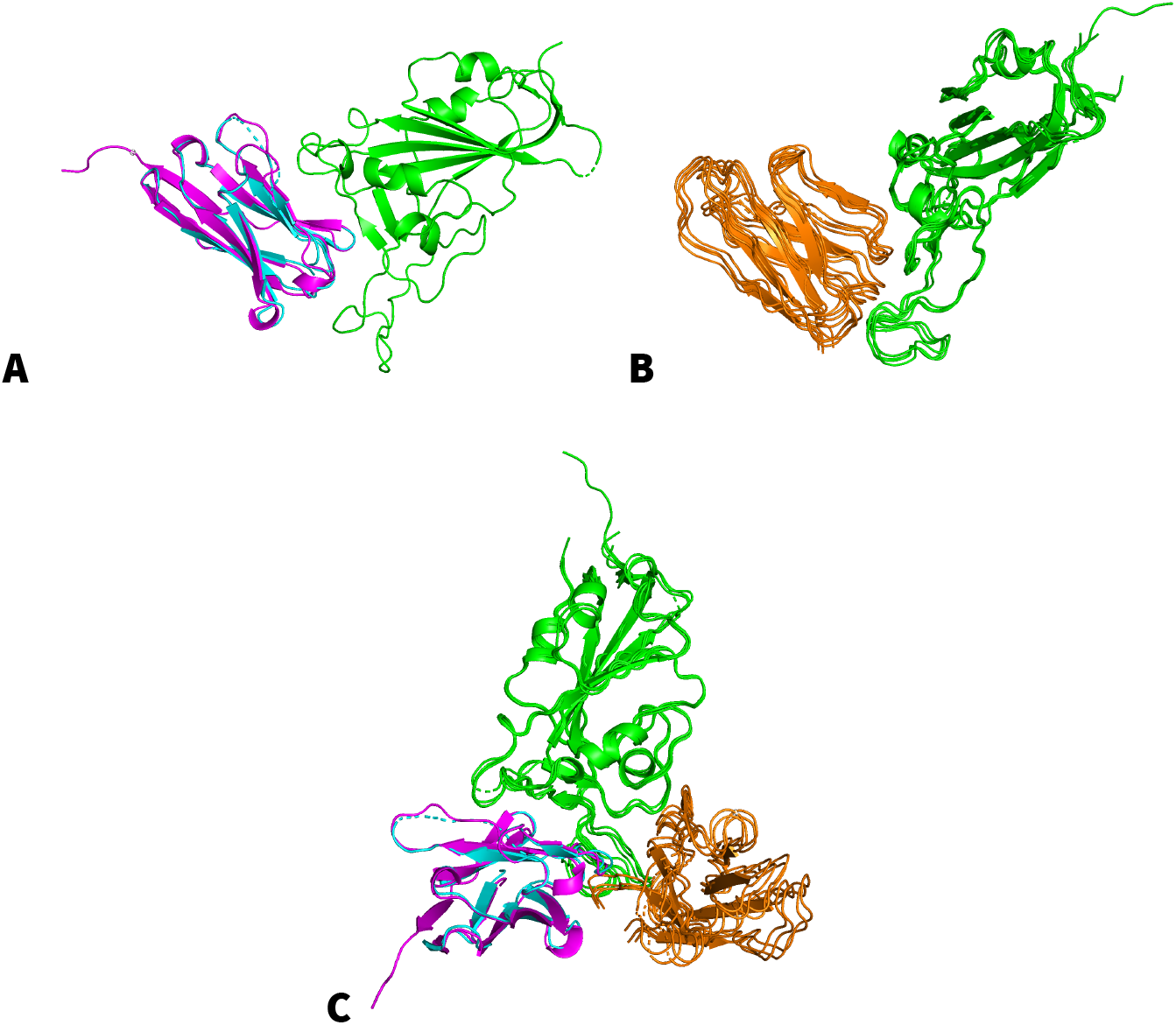
Putative binding modes of SARS-CoV2 receptor-binding domain (green) and (**A**) the top-ranking model of myelin-oligodendrocyte glycoprotein (cyan) with the homologue template of PDB entry 7C8V (pink) and (**B**) a cluster of secondary models (orange). **(C**) Display of both binding modes in complex with SARS-CoV2 receptor-binding domain.

**Table 3.** Multimeric frameworks identified for myelin-oligodendrocyte glycoprotein. Top scoring protein complex templates for the interaction between SARS-CoV2 spike protein (S) and myelin-oligodendrocyte glycoprotein receptor domain (MOG).

| PDB entry | Reference | Name | SM | TM | $E_{\text{contact}}$ | $Z_{\text{com}}$ |
| --- | --- | --- | --- | --- | --- | --- |
| 7C8V | Li, 2020 | Sybody SR4 | 0.90 | 0.82 | -8.57 | 45.8 |
| 6XC2 | Yuan, 2020 | Neutralizing Antibody CC12.1 | 0.86 | 0.83 | -3.53 | 55.2 |
| 7C01 | Shi, 2020 | Neutralizing Antibody | 0.86 | 0.83 | -3.28 | 55.3 |
| 7BZ5 | Wu, 2020 | Neutralizing Antibody | 0.85 | 0.83 | -2.79 | 55.3 |
| 6XC4 | Yuan, 2020 | Neutralizing Antibody CC12.3 | 0.84 | 0.81 | -2.54 | 53.4 |

The MOG receptor is associated with MOG antibody disease (MOGAD), a neuro-inflammatory condition that may cause inflammation of the optic nerve, the spinal cord and brain. Recent research has shown that SARS-CoV2 does trigger a relapse of MOGAD (Woodhall, 2020).

### SARS-CoV2 Spike protein (S) and Dipeptidyl Peptidase-4

High-scoring models were also generated for dipeptidyl peptidase-4 (DPP4, see **Fig. 5**), confirming the computational modeling results presented by Li et al. 2020. DPP4 is a cell surface glycoprotein receptor involved in T-cell activation and assumed to play a role in cell adhesion, migration and tube formation (Durinx, 2000).

**Figure 5.**
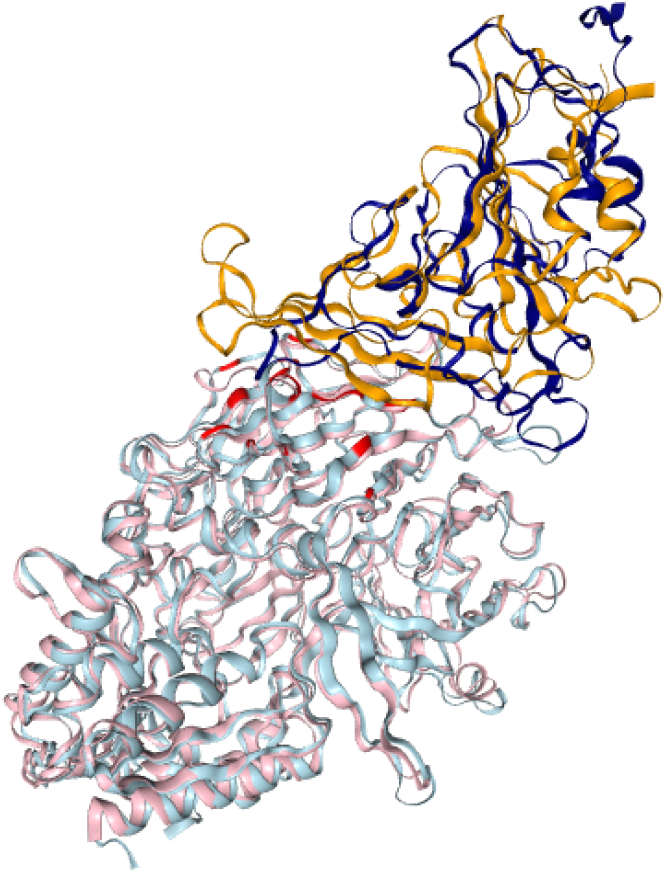
Putative binding mode between SARS-CoV2 (S) receptor-binding domain (RBD) (navy) and dipeptidyl peptidase-4 (DPP4) (cyan) with known MERS-CoV and DPP4 critical binding sites highlighted (red). The multimeric template framework is PDB entry 4L72 (pink), with the MERS-CoV receptor-binding domain (RBD) (orange).

Additionally, inhibiting DPP4 prevents glucagon release while increasing insulin secretion to decrease blood glucose levels (McIntosh, 2005). DPP4 is known to interact with MERS-CoV (Wang, 2013). The highest scoring template frameworks for DPP4 were PDB entry 4KR0 (Lu, 2013) with a Z_com_ score of 216.30 and PDB entry 4L72 (Wang, 2013) with 213.4. The resulting dimeric models are very similar to each other. The multimeric template matched the individual models with a TM-score of 0.62, a mean contact energy of - 6.7 and SM-score of 0.69.

Several sites are known to significantly contribute to the interaction between DPP4 and MERS-CoV’s receptor-binding domain (RBD). These are DPP4 residues K267, R336, R317, and Q344 (Song, 2014, Li, 2020, see **Fig. 5**) along with polymorphic sites as outlined in **Table 4**. Our method illustrates that SARS-Cov2’s S protein interacts with sites on DPP4 shared by MERS-CoV in addition to novel interaction sites (see **Table 4**). Additionally, half of fourteen critical binding sites (Letko, 2018) have been identified as polymorphic in Humans. Taken together, binding propensities between SARS-Cov2’s S protein and DPP4 might vary based on the population.

**Table 4.** shows sites on DPP4 that are critical in binding MERS-CoV’s receptor-binding domain (RBD) and sites predicted to interact with SARS-CoV2’s RBD. Sites on DPP4 that are known to be polymorphic in the Human population are highlighted. The results indicate that SARS-CoV2 interacts with sites on DPP4 known to be critical and additional novel sites.

| Sites on DPP4 | Interacting Virus | Critical, Polymorphic or Novel |
| --- | --- | --- |
| K267 | MERS-CoV | Critical and Polymorphic |
| R317 | MERS-CoV | Critical and Polymorphic |
| R336 | MERS-CoV | Critical |
| Q344 | MERS-CoV | Critical |
| T186 | SARS-CoV2 | Novel |
| T282 | SARS-CoV2 | Novel |
| S334 | SARS-CoV2 | Novel |

## Conclusion

Accurate identification of protein-protein interactions is essential to decipher cellular processes and detect novel drug targets. In the present work we implemented a Galaxy pipeline using the SPRING method which detects and structurally models protein-protein interactions by identifying distantly related protein complex structures with similar protein-protein interfaces.

The presented pipeline yields insights into the biochemical activity of SARS-CoV2 by identifying distant homologues with similar binding interfaces to Human proteins. For the papain-like protease of the non-structural protein 3 (Nsp3), we detected several ubiquitin-like substrates of which some have been experimentally confirmed. The method produced a top-ranking model for SARS-CoV’s spike protein (S) and dipeptidyl peptidase-4 (DPP4) in alignment with existing literature. Our method produced novel complex models between the S protein and myelin-oligodendrocyte glycoprotein (MOG). Here two top-ranking binding modes were produced. Experimental exploration will be needed to determine what impact these novel binding sites might play in pathogenicity, immune evasion, and adaptation. The prediction confidence relies on the accuracy of the homology match between templates, the structural fit and a knowledge-based contact potential, providing likely binding modes and interaction partners for further investigation. Only additional experimental validation can determine which or if any of the predicted binding modes occur in nature.

A limitation of our method is that it may produce high-confidence models between proteins which are localized in different subcellular regions. Existing literature has shown that such cross-interactions occur in a significant number of cases. In the present work we avoid filtering predicted protein interaction pairs by their corresponding subcellular locations since this would bias the obtained Matthew’s correlation coefficients. Identifying an accurate set of truly non-interacting protein pairs is critical and particularly challenging for the evaluation of protein-protein interactions. Randomly sampling protein pairs across a genome may lead to the inclusion of interacting protein pairs. A more accurate method is to sample non-interacting sets by pairing proteins from different subcellular regions as presented here. Yet another common suggestion is to exclude homologue protein pairs from the non-interacting set all together in order to avoid the inclusion of interacting pairs. This however is not an option due to the nature of the presented method which relies on homology detection to predict protein-protein interactions.

Another limitation is that homology modeling does rely on experimental templates. All of the fifteen most confident models derived for Human proteins interacting with Nsp3’s protease rely on four crystallographic complex structures.

This pipeline demonstrates the ability to detect interactome networks for a range of organisms. The increasing number of resolved co-crystal structures in the PDB, will continually improve the model quality and coverage over time (Chandonia and Brenner, 2006). Since the pipeline includes all data preparation steps no manual adjustment is required once new data has been published to the PDB. Galaxy enables users to employ the pipeline within their own methodologies and add or modify steps as required using Galaxy’s web-based workflow editor. Users are now able to reproduce and share the resulting interactome networks. The present contribution expands the repertoire of Galaxy tools to structural modeling methodologies, making them available for a large number of users. Recent advances in protein structure prediction and modeling (Kryshtafovych, 2019) complements existing sequence analysis tools and provides novel targets for drug discovery and elucidating biochemical processes through structural insights.

## Acknowledgments

The authors are grateful to the broader Galaxy community for their support and software development efforts. Thanks to Jessica Hamby for proofreading the manuscript.

## References

Afgan, Enis, et al. “The Galaxy Platform for Accessible, Reproducible and Collaborative Biomedical Analyses: 2018 Update.” Nucleic Acids Research, vol. 46, no. W1, 2018, doi:10.1093/nar/gky379.

Altschul, S. “Gapped BLAST and PSI-BLAST: a New Generation of Protein Database Search Programs.” Nucleic Acids Research, vol. 25, no. 17, 1997, pp. 3389–3402., doi:10.1093/nar/25.17.3389.

Benson, Dennis A., et al. “GenBank.” Nucleic Acids Research, vol. 43, no. D1, 2014, doi:10.1093/nar/gku1216.

Berman, H. M. “The Protein Data Bank.” Nucleic Acids Research, vol. 28, no. 1, 2000, pp. 235–242., doi:10.1093/nar/28.1.235.

Chandonia, John-Marc, and Steven E. Brenner. “The Impact of Structural Genomics: Expectations and Outcomes.” Science, vol. 311, no. 5759, 2006, pp. 347–351., doi:10.1126/science.1121018.

Daczkowski, Courtney M., et al. “Structurally Guided Removal of DeISGylase Biochemical Activity from Papain-Like Protease Originating from Middle East Respiratory Syndrome Coronavirus.” Journal of Virology, vol. 91, no. 23, 2017, doi:10.1128/jvi.01067-17.

Durinx, Christine, et al. “Molecular Characterization of Dipeptidyl Peptidase Activity in Serum.” European Journal of Biochemistry, vol. 267, no. 17, 2000, pp. 5608–5613., doi:10.1046/j.1432-1327.2000.01634.x.

Eshaghi, Majid, et al. “Rational Structure-Based Design of Bright GFP-Based Complexes with Tunable Dimerization.” Angewandte Chemie, vol. 127, no. 47, 2015, pp. 14158–14162., doi:10.1002/ange.201506686.

Gong, Weikang, et al. “Integrating Multimeric Threading with High-Throughput Experiments for Structural Interactome of Escherichia Coli.” 2020, doi:10.1101/2020.10.17.343962.

Guerler, Aysam, et al. “Mapping Monomeric Threading to Protein–Protein Structure Prediction.” Journal of Chemical Information and Modeling, vol. 53, no. 3, 2013, pp. 717–725., doi:10.1021/ci300579r.

Hunter, John D. “Matplotlib: A 2D Graphics Environment.” Computing in Science & Engineering, vol. 9, no. 3, 2007, pp. 90–95., doi:10.1109/mcse.2007.55.

Kezuka, Takeshi, and Hitoshi Ishikawa. “Diagnosis and Treatment of Anti-Myelin Oligodendrocyte Glycoprotein Antibody Positive Optic Neuritis.” Japanese Journal of Ophthalmology, vol. 62, no. 2, 2018, pp. 101–108., doi:10.1007/s10384-018-0561-1.

Kryshtafovych, Andriy, et al. “Critical Assessment of Methods of Protein Structure Prediction (CASP)—Round XIII.” Proteins: Structure, Function, and Bioinformatics, vol. 87, no. 12, 2019, pp. 1011–1020., doi:10.1002/prot.25823.

Lei, Jian, and Rolf Hilgenfeld. “Structural and Mutational Analysis of the Interaction between the Middle-East Respiratory Syndrome Coronavirus (MERS-CoV) Papain-like Protease and Human Ubiquitin.” Virologica Sinica, vol. 31, no. 4, 2016, pp. 288–299., doi:10.1007/s12250-016-3742-4.

Letko, Michael, et al. “Adaptive Evolution of MERS-CoV to Species Variation in DPP4.” Cell Reports, vol. 24, no. 7, 2018, pp. 1730–1737., doi:10.1016/j.celrep.2018.07.045.

Li, Dianfan, et al. “A Potent Synthetic Nanobody Targets RBD and Protects Mice from SARS-CoV-2 Infection.” 2020, doi:10.21203/rs.3.rs-75540/v1.

Li, Hong-Dong, et al. “A Network of Splice Isoforms for the Mouse.” Scientific Reports, vol. 6, no. 1, 2016, doi:10.1038/srep24507.

Li, Yu, et al. “The MERS-CoV Receptor DPP4 as a Candidate Binding Target of the SARS-CoV-2 Spike.” IScience, vol. 23, no. 8, 2020, p. 101400., doi:10.1016/j.isci.2020.101400.

Lu, Guangwen, et al. “Molecular Basis of Binding between Novel Human Coronavirus MERS-CoV and Its Receptor CD26.” Nature, vol. 500, no. 7461, 2013, pp. 227–231., doi:10.1038/nature12328.

Mcintosh, Christopher H.s., et al. “Dipeptidyl Peptidase IV Inhibitors: How Do They Work as New Antidiabetic Agents?” Regulatory Peptides, vol. 128, no. 2, 2005, pp. 159–165., doi:10.1016/j.regpep.2004.06.001.

Mering, C. Von. “STRING: Known and Predicted Protein-Protein Associations, Integrated and Transferred across Organisms.” Nucleic Acids Research, vol. 33, no. Database issue, 2004, doi:10.1093/nar/gki005.

Mesecar, A.d., and Y. Chen. “X-Ray Structure of MHV PLP2 (Cys1716Ser) Catalytic Mutant in Complex with Free Ubiquitin.” 2018, doi:10.2210/pdb5wfi/pdb.

Mesleh, Michael F., et al. “Marmoset Fine B Cell and T Cell Epitope Specificities Mapped onto a Homology Model of the Extracellular Domain of Human Myelin Oligodendrocyte Glycoprotein.” Neurobiology of Disease, vol. 9, no. 2, 2002, pp. 160–172., doi:10.1006/nbdi.2001.0474.

Naqvi, Ahmad Abu Turab, et al. “Insights into SARS-CoV-2 Genome, Structure, Evolution, Pathogenesis and Therapies: Structural Genomics Approach.” Biochimica Et Biophysica Acta (BBA) - Molecular Basis of Disease, vol. 1866, no. 10, 2020, p. 165878., doi:10.1016/j.bbadis.2020.165878.

Oughtred, Rose, et al. “The BioGRID Database: A Comprehensive Biomedical Resource of Curated Protein, Genetic, and Chemical Interactions.” Protein Science, vol. 30, no. 1, 2020, pp. 187–200., doi:10.1002/pro.3978.

Prescott, Lucas. “SARS-CoV-2 3CLpro Whole Human Proteome Cleavage Prediction and Enrichment/Depletion Analysis.” 2020, doi:10.1101/2020.08.24.265645.

Rao, V. Srinivasa, et al. “Protein-Protein Interaction Detection: Methods and Analysis.” International Journal of Proteomics, vol. 2014, 2014, pp. 1–12., doi:10.1155/2014/147648.

Shi, R., et al. “Molecular Basis for a Potent Human Neutralizing Antibody Targeting SARS-CoV-2 RBD.” 2020, doi:10.2210/pdb7c01/pdb.

Shin, Donghyuk, et al. “Papain-like Protease Regulates SARS-CoV-2 Viral Spread and Innate Immunity.” Nature, vol. 587, no. 7835, 2020, pp. 657–662., doi:10.1038/s41586-020-2601-5.

Shoemaker, Benjamin A, and Anna R Panchenko. “Deciphering Protein–Protein Interactions. Part I. Experimental Techniques and Databases.” PLoS Computational Biology, vol. 3, no. 3, 2007, doi:10.1371/journal.pcbi.0030042.

Song, Wenfei, et al. “Identification of Residues on Human Receptor DPP4 Critical for MERS-CoV Binding and Entry.” Virology, vol. 471-473, 2014, pp. 49–53., doi:10.1016/j.virol.2014.10.006.

Steinegger, Martin, et al. “HH-suite3 for Fast Remote Homology Detection and Deep Protein Annotation.” 2019, doi:10.1101/560029.

Uniprot Consortium, The. “UniProt: the Universal Protein Knowledgebase.” Nucleic Acids Research, vol. 46, no. 5, 2018, pp. 2699–2699., doi:10.1093/nar/gky092.

Valencia, Inés, et al. “DPP4 And ACE2 in Diabetes and COVID-19: Therapeutic Targets for Cardiovascular Complications?” Frontiers in Pharmacology, vol. 11, 2020, doi:10.3389/fphar.2020.01161.

Vangaveti, Sweta, et al. “Integrating Ab Initio and Template-Based Algorithms for Protein–Protein Complex Structure Prediction.” Bioinformatics, vol. 36, no. 3, 2019, pp. 751–757., doi:10.1093/bioinformatics/btz623.

Wang, Nianshuang, et al. “Structure of MERS-CoV Spike Receptor-Binding Domain Complexed with Human Receptor DPP4.” Cell Research, vol. 23, no. 8, 2013, pp. 986–993., doi:10.1038/cr.2013.92.

Wells, James A., and Christopher L. Mcclendon. “Reaching for High-Hanging Fruit in Drug Discovery at Protein–Protein Interfaces.” Nature, vol. 450, no. 7172, 2007, pp. 1001–1009., doi:10.1038/nature06526.

Woodhall, Mark, et al. “Case Report: Myelin Oligodendrocyte Glycoprotein Antibody-Associated Relapse With COVID-19.” Frontiers in Neurology, vol. 11, 2020, doi:10.3389/fneur.2020.598531.

Wu, Yan, et al. “A Non-Competing Pair of Human Neutralizing Antibodies Block COVID-19 Virus Binding to Its Receptor ACE2.” 2020, doi:10.1101/2020.05.01.20077743.

Yuan, Meng, et al. “Structural Basis of a Shared Antibody Response to SARS-CoV-2.” Science, vol. 369, no. 6507, 2020, pp. 1119–1123., doi:10.1126/science.abd2321.

Zhang, Chengxin, et al. “Functions of Essential Genes and a Scale-Free Protein Interaction Network Revealed by Structure-Based Function and Interaction Prediction for a Minimal Genome.” Journal of Proteome Research, vol. 20, no. 2, 2021, pp. 1178–1189., doi:10.1021/acs.jproteome.0c00359.

Zhang, Jian, and Yang Zhang. “A Novel Side-Chain Orientation Dependent Potential Derived from Random-Walk Reference State for Protein Fold Selection and Structure Prediction.” PLoS ONE, vol. 5, no. 10, 2010, doi:10.1371/journal.pone.0015386.

Zhang, Y. “TM-Align: a Protein Structure Alignment Algorithm Based on the TM-Score.” Nucleic Acids Research, vol. 33, no. 7, 2005, pp. 2302–2309., doi:10.1093/nar/gki524.

Zhou, Mi, et al. “ChemInform Abstract: Current Experimental Methods for Characterizing Protein-Protein Interactions.” ChemInform, vol. 47, no. 24, 2016, doi:10.1002/chin.201624266.

Zhou, Peng, et al. “A Pneumonia Outbreak Associated with a New Coronavirus of Probable Bat Origin.” Nature, vol. 579, no. 7798, 2020, pp. 270–273., doi:10.1038/s41586-020-2012-7.

Zhou, Siwei, et al. “Myelin Oligodendrocyte Glycoprotein Antibody–Associated Optic Neuritis and Myelitis in COVID-19.” Journal of Neuro-Ophthalmology, vol. 40, no. 3, 2020, pp. 398–402., doi:10.1097/wno.0000000000001049.

